# High-dimensional multiomics reveals perturbations to IL-6/IL-6R axis and RUNX3 in CD4^+^ T cells during third trimester pregnancy

**DOI:** 10.64898/2026.03.26.711478

**Authors:** Jennifer R Habel, Thi H O Nguyen, Natasha de Alwis, E Kaitlynn Allen, Shihan Li, Morgan J Skinner, Jennifer A Juno, Stephen J Kent, Katherine Bond, Deborah A Williamson, Martha Lappas, Natalie J Hannan, Susan Walker, Jan Schroeder, Jeremy Chase Crawford, Paul G Thomas, Katherine Kedzierska, Louise C Rowntree

**Affiliations:** Department of Microbiology and Immunology, University of Melbourne, at the Peter Doherty Institute for Infection and Immunity, Melbourne, Victoria, Australia; The Department of Obstetrics, Gynaecology and Newborn Health, University of Melbourne, Heidelberg, Victoria, Australia; Mercy Perinatal, Mercy Hospital for Women, Heidelberg, Victoria, Australia; Department of Immunology, St. Jude Children’s Research Hospital, Memphis, Tennessee 38105, USA; Computational Sciences Initiative, Department of Immunology and Microbiology, University of Melbourne at the Peter Doherty Institute for Infection and Immunity, Melbourne, Australia; Victorian Infectious Diseases Reference Laboratory, at the Peter Doherty Institute for Infection and Immunity, Melbourne, Victoria, Australia; Department of Microbiology, Royal Melbourne Hospital, Melbourne, Victoria, Australia; Victorian Infectious Diseases Service, Royal Melbourne Hospital, at the Peter Doherty Institute for Infection and Immunity, Melbourne, Victoria, Australia; Department of Infectious Diseases, The University of Melbourne at the Peter Doherty Institute for Infection and Immunity, Melbourne, Victoria, Australia; School of Medicine, University of St Andrews, Scotland KY16 9TF, United Kingdom; Department of Host-Microbe Interactions, St. Jude Children’s Research Hospital, Memphis, Tennessee 38105, USA; Immunology and Vaccine Development Program, Vaccine and Infectious Disease Division, Fred Hutchinson Cancer Center, Seattle 98109, Washington, USA

**Author notes:** authors contributed equally (Katherine Kedzierska, Louise C Rowntree). Ethics statement: Experiments conformed to the Declaration of Helsinki Principles and the Australian National Health and Medical Research Council Code of Practice. Written informed consents were obtained from all participants prior to the study. The studies were approved by the Mercy Health (R14/25 and R04/29), Melbourne Health (HREC/66341/MH-2020), and University of Melbourne (#1443540, 2024-13344-58055-11, 2020-20782-12450-1) human research ethics committees (HRECS).

**Keywords:** CD4^+^ T cell, pregnancy, human, IL-6R

## Abstract

**Objectives:** CD4^+^ T cells play key roles in regulating immune responses during pregnancy, therefore we aimed to understand the CD4^+^ T cell surface proteome and transcriptome during pregnancy.

**Methods:** CD4^+^ T cells were analysed in blood and decidua from term-pregnancies (>37 weeks), and non-pregnant blood. >350 surface proteins were screened via flow cytometry, and transcriptomes were analysed using single-cell RNA sequencing with >130 CITE-seq barcoded antibodies.

**Results:** Surface protein screening identified changes to ILT4/CD85d, CD9, IFN-γ receptor β-chain, CX3CR1 and CCR5 in the pregnant blood and decidual CD4^+^ T cells. CX3CR1 and CCR5 had the highest expression on the effector-memory T cell (T_EM_) subset in the blood, with expression consistent across subsets in decidua. CD126/IL-6R was lower in pregnant blood and decidual CD4^+^ T cells, while scRNAseq identified enrichment in the IL-6R signalling pathway in naive CD4^+^ T cells in pregnant blood. Both sIL-6R and IL-6 concentrations were increased in plasma during pregnancy, suggesting perturbations to the IL-6/IL-6R signalling axis. Meanwhile, decidual CD4^+^ T cells had increased expression of transcription factor RUNX3 in the CD69^+^ tissue-resident-like subset.

**Conclusions:** Our findings demonstrate altered molecular expression in CD4^+^ T cells during pregnancy. This provides important mechanistic insight of their adaptation and regulation during placental development, which may drive placental dysfunction or pregnancy complications including preeclampsia, fetal growth restriction and stillbirth. These new data may inform future studies that focus on determining the significance of differentially-expressed immune features in pregnancy to identify potential targets for immune modulation to treat pregnancy complications and infections.

## INTRODUCTION

During pregnancy maternal CD4^+^ T cells play a vital role in maintaining tolerance ^1^. There are dynamic changes to the frequencies of regulatory CD4^+^ T cells (T_reg_) across pregnancy trimesters^2^. Within the decidua CD4^+^ T cells control inflammation caused by blastocyst implantation^3^. Impairments to decidual CD4^+^ T cells have been implicated in obstetric complications, including preeclampsia, recurrent pregnancy loss, and spontaneous preterm labour. While the exact mechanisms underlying CD4^+^ T cell involvement in pregnancy complications are unknown, hypotheses suggest an imbalance in the Th1/Th2/Th17/T_reg_ subsets^4^. Furthermore, reduced numbers of decidual regulatory CD25^bright^CD4^+^ T cells is associated with spontaneous abortion, exemplifying the importance of CD4^+^ T cells in healthy pregnancy^5,6^. In fact pregnancy-acquired CD4^+^ T_reg_ have been shown to improve future pregnancy outcomes in mouse models, suggesting that pregnancy has long-lasting effects on CD4^+^ T cell functionality^7^.

In addition to CD4^+^ T cells being central to immune regulation in pregnancy, they play key roles in antiviral immunity. Their functions include providing help to B cells, immunoregulation, and cytotoxicity^8-10^. As CD4^+^ T cells undergo changes during pregnancy, this could impact the ability to fight viral infections, which is of particular importance in the third pregnancy trimester when the risk of severe infection outcomes is highest. Therefore, our study used high-dimensional techniques to investigate differential gene and protein expression on CD4^+^ T cells between full-term 3^rd^ trimester pregnant and non-pregnant women, with a focus of identifying potential targets that have implications in antiviral immunity. We screened >350 surface proteins on naïve, memory, and regulatory-like CD25^+^ CD4^+^ T cells and performed single-cell RNA sequencing (scRNAseq) using 130 surface CITE-seq antibodies to measure whole transcriptomes and surface proteomes to provide insights on the altered phenotypes during pregnancy and their implications for antiviral immunity.

## RESULTS AND DISCUSSION

### Pregnancy changes CD4^+^ T cell surface protein expression

Given the importance of CD4^+^ T cells during pregnancy, and our current understanding of their critical importance in placental establishment and function^11-13^, we analysed the proportions of memory and regulatory subsets in the blood and decidua (Fig 1A). Compared to non-pregnant blood, there were higher proportions of the CD27^+^CD45RA^-^ central memory-like (T_CM_) subset in the pregnant blood (Fig 1B). Meanwhile, increased frequencies of CD27^-^CD45RA^-^ effector memory-like (T_EM_) CD4^+^ T cells were detected in the decidua compared to matched pregnant blood (Fig 1B). The frequencies of regulatory CD4^+^ T cells (Treg) were defined by CD25 expression, with similar frequencies observed between pregnant and non-pregnant blood as well as pregnant blood and decidua (Fig 1C-D). Most CD4^+^ T cells across all groups were T_Naive_ (mean ∼42-62%) or T_CM_ (mean ∼32-45%), while the CD25^+^ T_reg_-like made up ∼10-15% of CD4^+^ T cells (Fig 1B,D). Given the low frequencies of the T_SCM_ and T_EMRA_ subsets, we focused on T_Naive_, T_CM_, T_EM,_ and T_reg_ subsets in the surface protein screening analysis.

**Figure 1.**
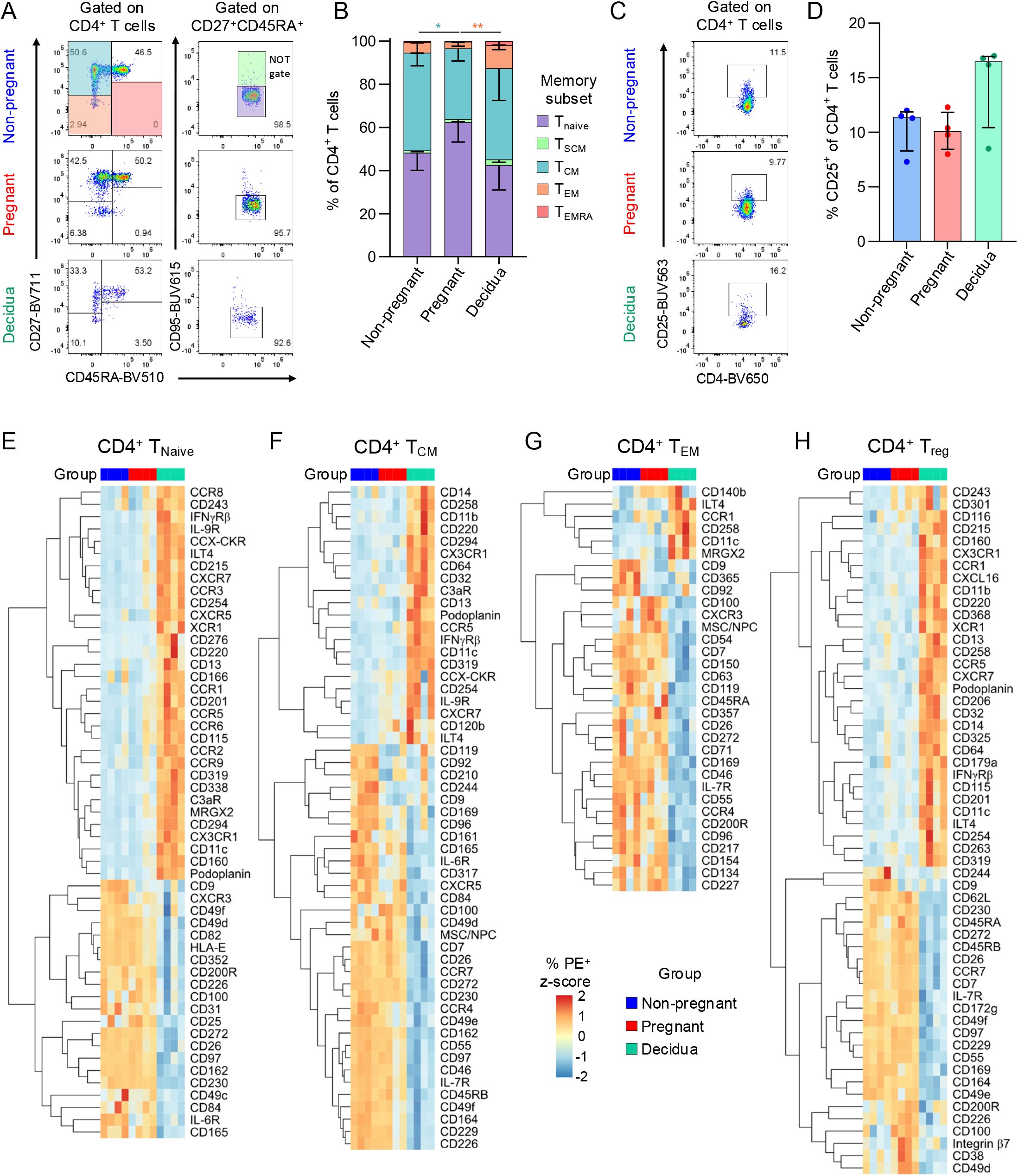
Screening CD4^+^ T cell surface protein expression. (A) Representative flow cytometry plots of staining for CD27, CD45RA, and CD95 to determine CD4^+^ T cell memory phenotypes in non-pregnant and pregnant blood, and decidua. (B) Frequencies of T_Naive_, T_SCM_, T_CM_, T_EM_ and T_EMRA_ CD4^+^ T cell subsets in non-pregnant (n=4) and pregnant blood (n=4), and decidua (n=4). (C) Representative flow cytometry plots of CD25 staining on CD4^+^ T cells to identify T_reg_ cells in non-pregnant and pregnant blood, and decidua. (D) Frequencies of CD25^+^ regulatory CD4^+^ T cells in non-pregnant (n=4) and pregnant blood (n=4), and decidua (n=4). (E-H) Heatmaps depicting significantly differentially expressed surface proteins in T_Naive_, T_CM_, T_EM_, and T_reg_ CD4^+^ T cell subsets in non-pregnant (n=4) and pregnant blood (n=4), and decidua (n=4). Markers with >10% PE^+^ in at least one sample were included in analysis, and those with zero or near zero variance across all samples were removed.

To determine differential protein expression across peripheral blood CD4^+^ T cell subsets during pregnancy, frequencies were compared to those of the non-pregnant group. Within the T_Naive_ subset, there was decreased expression of CD9 and IL-6R (CD126) during pregnancy (Fig 1E; Supplementary Table 1). CD9 and IL-6R were also decreased in the T_CM_ subset in the pregnant group, while CD85d was increased although at a lower magnitude (Fig 1F; Supplementary Table 2). Within both the T_EM_ and T_reg_ subsets, markers with increased frequencies during pregnancy included CD100, CD200R, and CD272 (BTLA), while CD9 was decreased (Fig 1G-H; Supplementary Tables 3-4).

We next compared decidual and pregnant peripheral blood CD4^+^ T cells to identify tissue-specific features. Within the T_Naive_, T_CM_, and T_reg_ subsets, chemokine receptors with the highest increase in expression included CCR5 and CX3CR1 (Fig 1E-H; Supplementary Tables 5-8). In contrast, CCR5 and CX3CR1 had similar levels of expression between the blood and decidua within the T_EM_ subset, which instead had lower frequencies of CCR4^+^ and CXCR3^+^ cells (Fig 1E-H). Across all subsets, except for T_CM_, CD85d (ILT4) was increased on decidual CD4^+^ T cells. ILT4 has been shown to increase on activated CD4^+^ T cells, and is a receptor for semaphorin 4A, which is produced by the fetal trophoblast^14,15^. The interaction of semaphorin 4A with ILT4 on CD4^+^ T cells is linked with Th2 differentiation^15^, which could have a role in CD4^+^ T cell regulation at the maternal-fetal interface.

### Linking differentially expressed proteins in pregnancy to CD4^+^ T cell subsets

We subsequently investigated whether differentially expressed markers were specific to each CD4^+^ T cell subset or were shared across multiple subsets within pregnant blood and decidua. CD9 was the only protein commonly downregulated across the four CD4^+^ T cell subsets in pregnant blood, while CD11c was one of three proteins increased in all decidual subsets (Fig 2A). CD9-expressing peripheral blood CD4^+^ T cells could be due to lymphocyte-platelet conjugates^16^, and the lower frequencies of CD9^+^ CD4^+^ T cells observed in pregnancy could be linked to the characteristic decline in platelet numbers during gestation^17^. While little is known about the role of CD11c^+^ CD4^+^ T cells in pregnancy, dysregulation of CD11c^+^ CD8^+^ T cells in the decidua is implicated in early pregnancy loss^18^. Our further analysis of surface protein expression across memory subsets focused on proteins with a high magnitude of differential expression and implications for CD4^+^ T cell functionality. Therefore, we further assessed expression of IFN-γ receptor β-chain, CD85d/ILT4, CD9, CD126/IL-6R, CCR5, and CX3CR1 across CD4^+^ T_Naive_, T_CM_, T_EM_, and T_reg_ subsets.

**Figure 2.**
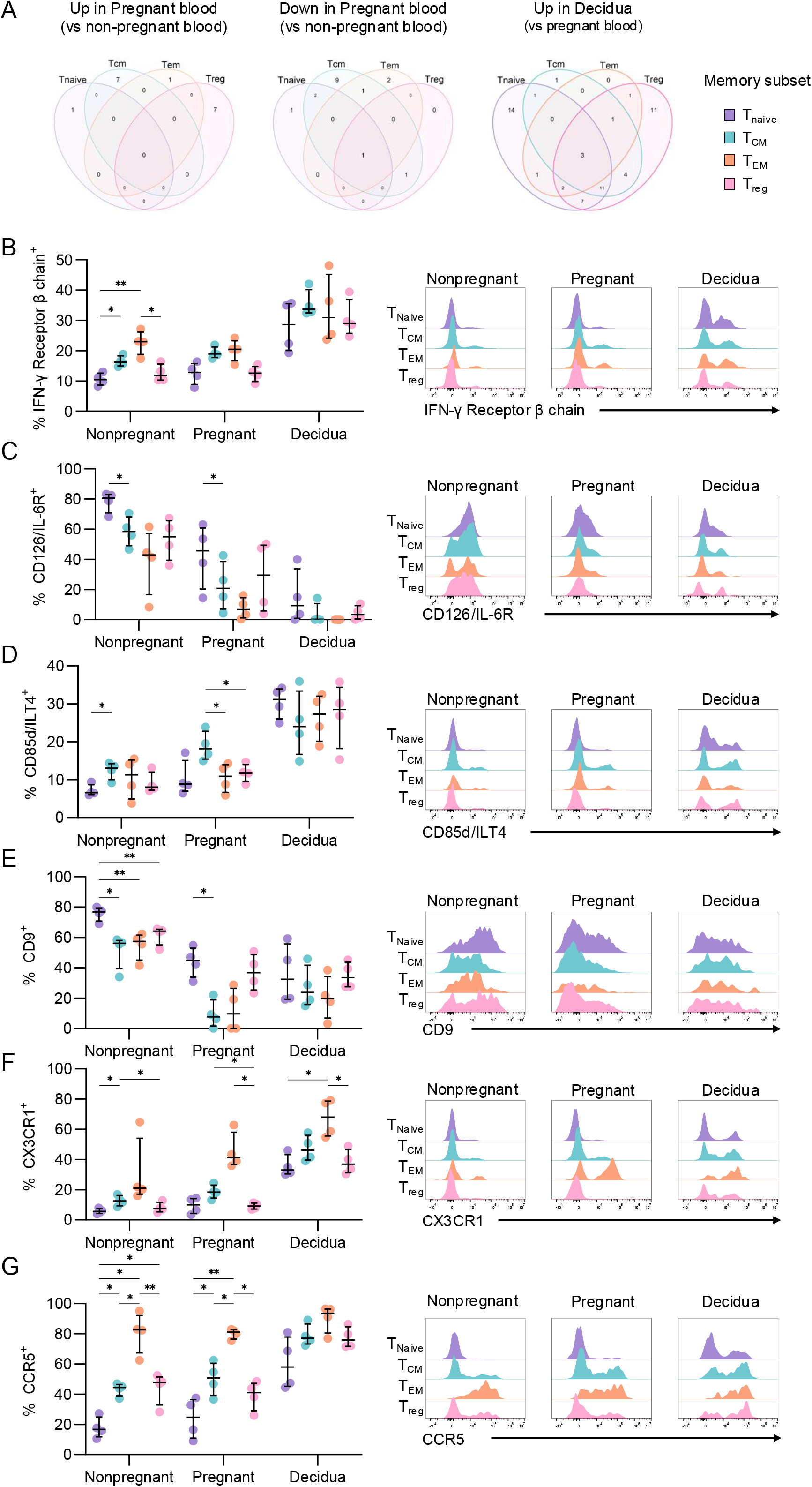
Linking differentially expressed surface proteins with CD4^+^ T cell subsets. (A) Venn diagrams depicting overlap in differential protein expression across T_Naive_, T_CM_, T_EM_, and T_reg_ subsets for proteins upregulated (left) or downregulated (middle) in pregnant blood compared to non-pregnant blood, and upregulated (right) in the decidua compared to pregnant blood. (B-G) Frequencies and representative histograms of flow cytometry data of selected proteins IFN-γ receptor β (B), CD126/IL-6R (C), CD85d/ILT4 (D), CD9 (E), CX3CR1 (F), and CCR5 (G) across T_Naive_, T_CM_, T_EM_, and T_reg_ subsets and non-pregnant (n=4) and pregnant blood (n=4), and decidua (n=4). Statistic shown is a 2-way ANOVA performed within the non-pregnant, pregnant, or decidua groups to determine differences across T cell subsets within each group. *p<0.05, **p<0.01, ***p<0.001.

IFN-γ receptor β-chain is a component of the IFN-γ receptor, which sensitizes CD4^+^ T cells to IFN-γ signalling. IFN-γ receptor β-chain had increasing expression across differentiation states in the non-pregnant group, with T_CM_ and T_EM_ being higher than T_Naive_, while T_reg_ had similar frequencies as T_Naive_ (Fig 2B). Meanwhile frequencies were similar across subsets within the pregnant blood and decidua groups.

IL-6 plays a significant role in CD4^+^ T cell activation and differentiation, and expression of its receptor, CD126/IL-6R, in part, dictates sensitivity to IL-6 signalling^19^. CD126/IL-6R was most highly expressed on T_Naive_ cells in the pregnant and non-pregnant blood, while expression in the decidua was consistently low across subsets (Fig 2C). The lower expression of IL-6R in the blood and decidua during pregnancy could be due to intrinsic downregulation thereby decreasing susceptibility to IL-6 signalling, or alternatively due to shedding of soluble IL-6R (sIL-6R), known to occur during CD4^+^ T cell activation^20^.

CD85d/ILT4 is an inhibitory receptor, however its expression is only upregulated upon CD4^+^ T cell activation^15^. CD85d/ILT4 had consistently high expression across all subsets in the decidua, while T_CM_ had the highest frequency of CD85d^+^ cells in the pregnant group (Fig 2D). Given that the pregnant group consists of term pregnancies, there could be increased inflammation leading into labour/parturition that activates CD4^+^ T cells, contributing to increased CD85d expression.

Frequencies of CD9^+^ cells were highest in T_Naive_ for both the pregnant and non-pregnant groups, while the decidua had consistent expression across all subsets (Fig 2E). This could indicate that the T_Naive_ subset is more prone to forming lymphocyte-platelet conjugates, however further studies are required to determine the molecular features behind this interaction and its functional significance.

Finally, CX3CR1 and CCR5 are involved in cell migration, with their ligands being CX3CL1 and CCL3/4/5, respectively. Across all groups, CX3CR1 expression was highest in the T_EM_ subset, while T_Naive_ and T_reg_ subsets had the lowest expression (Fig 2F). This is in line with a recent study which showed CX3CR1 increases with T cell differentiation, with the highest expression observed in the terminally-differentiated subset^21^. Similarly, CCR5 expression was highest on T_EM_ within pregnant and non-pregnant blood, while frequencies were similar across all subsets in the decidua (Fig 2G). The ligands for CCR5, CCL3 and CCL5, are produced by the fetal trophoblast, likely contributing to recruitment of CCR5^+^ CD4^+^ T cells to the decidua.

### Pregnancy alters CD4^+^ T_Naive_ and T_CM_ transcriptomes

To determine underlying transcriptomic differences that could potentially impact CD4^+^ T cell surface protein expression, we performed single-cell RNA sequencing on peripheral blood CD4^+^ T cells from pregnant and non-pregnant individuals, and pregnancy-matched decidua. Dimensionality reduction resulted in subsets of T_Naïve_ and T_CM_ CD4^+^ T cells based on protein expression of CD27, CD45RA/RO, and CD95, and RNA expression of *CCR7* and *FAS* (Fig 1A-B). Based on this, UMAP clusters 0, 2 and 4 were annotated as T_Naive_ and clusters 1, 3 and 5 as T_CM_. Peripheral blood CD4^+^ T cells were observed across all UMAP clusters, with cluster 4 and 5 uniquely being made up of CD4^+^ T cells from pregnant individuals, while decidual CD4^+^ T cells were predominantly observed in clusters 2 and 3 (Fig 3A-D).

**Figure 3.**
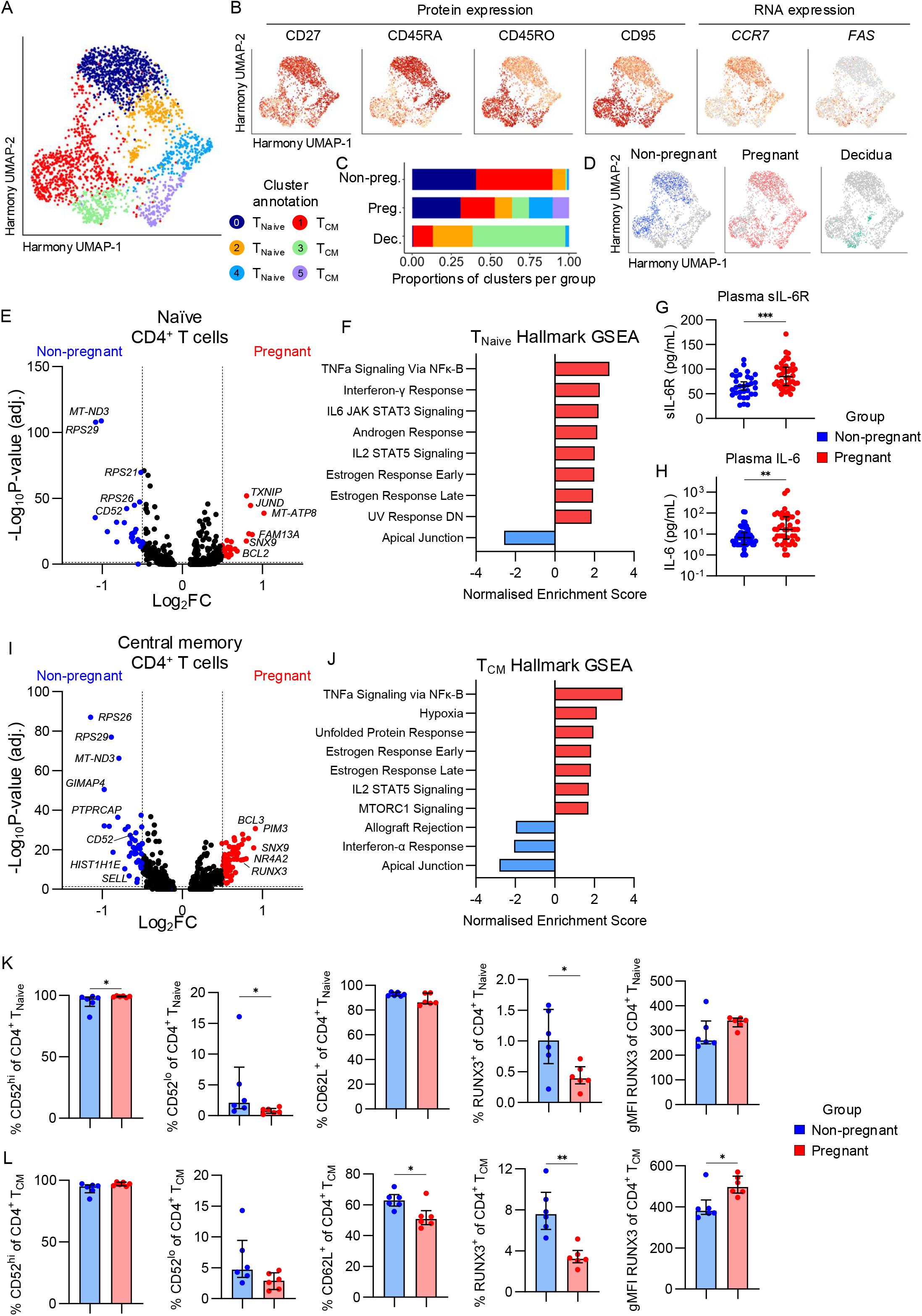
Defining CD4^+^ T cell transcriptomes during pregnancy with single-cell RNA sequencing. (A) Harmony UMAP of CD4^+^ T cells depicting 6 clusters. Non-pregnant n=3, Pregnant n=4, Decidua n=3. (B) Harmony UMAPs depicting protein expression via CITE-seq of CD27, CD45RA, CD45RO, and CD95, and RNA expression of *CCR7* and *FAS* (encoding CD95) used to define CD4^+^ T cell subsets. Non-pregnant n=3, Pregnant n=4, Decidua n=3. (C) Proportions of cells in each Harmony UMAP cluster for non-pregnant (n=3), pregnant (n=4), and decidua (n=3) groups. (D) Harmony UMAPs highlighting cells from the non- pregnant (n=3), pregnant (n=4), and decidua (n=3) groups. (E) Volcano plot showing differential gene expression between pregnant (n=4) and non-pregnant (n=3) T_Naive_ CD4^+^ T cells. (F) Gene set enrichment analysis showing the Normalised Enrichment Scores for all significantly enriched Hallmark gene sets in the T_Naive_ subset. (G-H) Concentration of sIL-6R and IL-6 in blood plasma from pregnant (n=44) and non-pregnant (n=47) individuals. IL-6 concentrations are derived from our previous COVID-19 pregnancy study^24^. (I) Volcano plot showing differential gene expression between pregnant (n=4) and non-pregnant (n=3) T_CM_ CD4^+^ T cells. (J) Gene set enrichment analysis showing the Normalised Enrichment Scores for all significantly enriched Hallmark gene sets in the T_CM_ subset. (K-L) Frequencies of CD52^hi^, CD52^lo^, CD62L^+^ and RUNX3^+^, and gMFI of RUNX3, for (K) T_Naive_ and (L) T_CM_ CD4^+^ T cell subsets in pregnant (n=6) and non-pregnant (n=6) groups. Statistic shown is a Mann- Whitney test. (E,I) Vertical dashed lines indicate a Log_2_ fold-change (Log_2_FC) of 0.5 for comparisons in the blood and 0.75 for comparison between blood and decidua, horizontal dashed line indicates the Log_10_ adjusted p-value cutoff for significance of 1.3. *p<0.05.

Differential gene expression analysis of the T_Naive_ subset in pregnant and non-pregnant peripheral blood showed increased expression of 25 genes (>0.5 log_2_FC) during pregnancy, with top 10 including *MT-ATP8, FAM13A, JUND, ANK3, TXNIP, SNX9, BCL2, GPRIN3, NFKBIA*, and *SOCS3* (Fig 3E; Supplementary Table 9). Meanwhile, 25 genes had higher expression in the non-pregnant group, including *MTRNR2L8, RPS29, MT-ND3, ACTB, CD52, HIST1H4C, LBH, RPS10, GIMAP4*, and *MTRNR2L12* (Fig 3E; Supplementary Table 9). Hallmark gene set enrichment analysis (GSEA) identified that T_Naive_ cells from pregnant women were enriched for genes involved in the IL-6/JAK/STAT3 and IL-2/STAT5 signalling pathways, indicating enhanced activation (Fig 3F). Our surface protein screening identified that T_Naive_ had the highest frequencies of IL-6R^+^ cells, with lower frequencies in pregnancy. Given that transcriptomics identified enhanced IL-6 mediated signalling, downregulation of IL-6R could be part of a negative-feedback loop, or molecules could be shed from the surface as sIL-6R resulting in reduced frequencies of IL-6R^+^ cells. To further investigate the IL-6 axis in pregnancy, we measured sIL-6R and IL-6 in the blood plasma of pregnant and non-pregnant individuals. Indeed, concentrations of both sIL-6R and IL-6 were higher in pregnancy, supporting that IL-6 mediated signalling is perturbed in during pregnancy (Fig 3G-H).

Within the T_CM_ subset, 59 genes were increased in pregnancy, including *PIM3, SNX9, CNOT6L, NR4A2, PDE4D, MT-ATP8, BCL3, PDE3B, RUNX3*, and *NINJ1*, while 43 genes were increased in the non-pregnant group including *MTRNR2L8, RPS26, GIMAP4, RPS29, PTPRCAP, MT-ND3, HIST1H1E, IL7R, RAC2*, and *SELL* (Fig 3G; Supplementary Table 10). Similar to T_Naive_, GSEA showed an enrichment for the Early and Late Estrogen Response, TNF signalling, and IL-2/STAT5 signalling in the T_CM_ during pregnancy (Fig 3F,H). Given that estrogen increases during pregnancy and can directly induce expression of anti-apoptotic proteins like BCL2, also found to be increased in the pregnant group, there might be enhanced control of CD4^+^ T cell death during pregnancy (Fig 3E,F).

To determine whether DEGs between pregnant and non-pregnant groups were different at the protein level, flow cytometry was performed to analyse CD52, CD62L and RUNX3 on the T_Naive_ and T_CM_ subsets (Supplementary Figure 2). In contrast to transcriptomics, CD52 expression remained high in both the pregnant and non-pregnant groups, suggesting the minor decrease in the non-pregnant group unlikely to be physiologically relevant (Fig 3K). However, reflecting the transcriptomic data, there was a lower frequency of CD62L^+^ T_CM_ and an increased RUNX3 gMFI in T_CM_ during pregnancy (Fig 3K-L). The frequency of RUNX3^+^ T_CM_ cells was higher in the non-pregnant group, suggesting that fewer T cells express RUNX3 but the amount is increased in pregnancy (Fig 3L).

Taken together, our surface protein screening and transcriptomic datasets identified consistent alterations to molecules involved in IL-6 and estrogen signalling in T_Naive_ and T_CM_ subsets during pregnancy, both of which upregulate expression of anti-apoptotic molecules like BCL2. Future studies investigating mechanisms of apoptotic regulation in T cells during pregnancy, and in pregnancy complications that are underpinned by a dysfunctional placenta (including preeclampsia and fetal growth restriction), will identify its importance and whether this axis is disrupted during pregnancy complications.

### RUNX3 involved in decidual tissue-resident CD4^+^ T cell phenotype

Further to the differences observed between pregnant and non-pregnant CD4^+^ T cells in the blood, we compared pregnant peripheral blood and decidual CD4^+^ T cells. Decidual CD4^+^ T cells displayed a more activated transcriptome, with upregulation of genes like *GZMK, PRF1*, and *ICOS*, in addition to genes involved in apoptosis and inflammatory signalling like *LMNA, ANXA1, MCL1, CD69, JUN, IFNGR1, TNF*, and *FAS* (Fig 4A; Supplementary Table 11). These genes contributed to the enrichment of Hallmark Gene Sets related to T cell activation, such as TNF signalling, Inflammatory Response, IL-2/STAT5 signalling, and IFN-γ response (Fig 4B). Further analysis by flow cytometry showed, in contrast to transcriptomics, granzyme K was higher in CD4^+^ T cells from pregnant blood than the decidua (Fig 4C). Meanwhile, decidual CD4^+^ T cells had higher CD69 expression, particularly within the T_CM_ subset (Fig 4D). Given RUNX3 is associated with tissue-residency in CD8^+^ T cells^22^, CD62L with central memory, and CD95 with T cell differentiation, we assessed these markers on decidual CD69^+^ and CD69^-^ CD4^+^ T cells. The CD69^+^ subset had significantly higher frequencies of RUNX3^+^ and CD95^+^ T cells, and lower frequencies of CD62L^+^ T cells (Fig 4E-F). This suggests RUNX3 could be important for decidual tissue-residency, and that the CD69^+^ CD4^+^ T cells in the decidua are more differentiated due to increased CD95 and decreased CD62L, in line with a previous report^23^.

**Figure 4.**
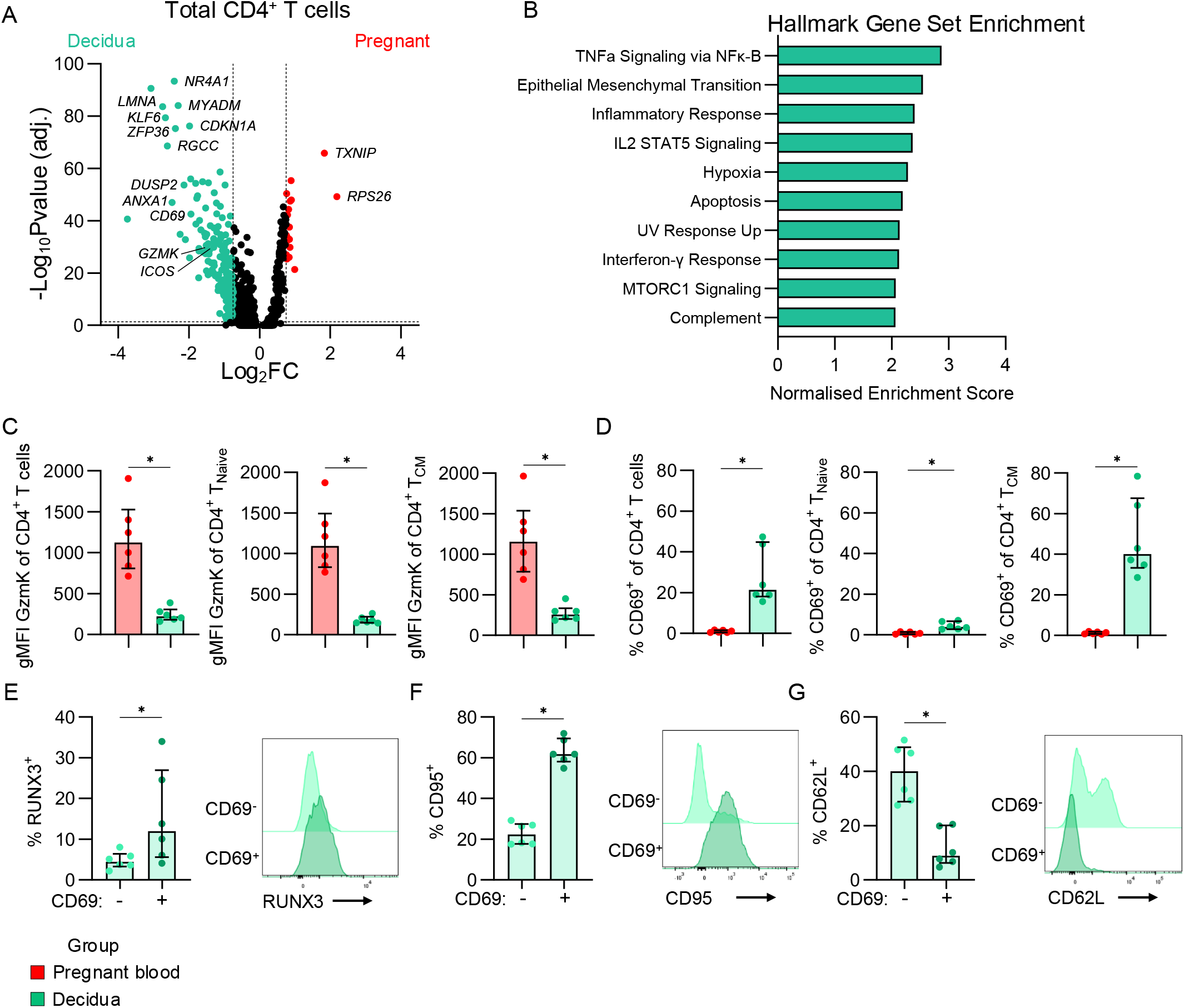
Defining tissue-resident CD4^+^ T cells in the decidua. (A) Volcano plot showing differential gene expression between pregnant (n=4) and decidual (n=3) total CD4^+^ T cells. (B) Gene set enrichment analysis showing the Normalised Enrichment Scores for the top 10 significantly enriched Hallmark gene sets in decidual CD4^+^ T cells. (C-D) gMFI of granzyme K (C) and frequencies of CD69^+^ (D) in total CD4^+^ T cell, T_Naive_ and T_CM_ subsets in the pregnant blood (n=6) and decidua (n=6). Statistic shown is a Wilcoxon test. *p<0.05. (E-G) Frequencies and representative flow cytometry histograms showing RUNX3 (E), CD95 (F), and CD62L (G) in the CD69^-^ and CD69^+^ subsets in the decidua (n=6).

### Limitations to the study

This study is an observational and correlative study used high-dimensional approaches to explore phenotypic and transcriptional differences to CD4^+^ T cell immunity in 3^rd^ trimester pregnancy. The interpretation of results is limited by the relatively small sample size. Due to the limited sample size, certain confounders could not be accounted for including gravidity, parity and fetal sex of the pregnant group. Additionally, prior pregnancy information for the non-pregnant group is unknown. Future expanded studies should account for these variables to identify if they contribute to the observed differences. Finally, the focus of this study was on 3^rd^ trimester pregnancies due to the increased risk of severe viral infections, therefore it is unknown whether these findings apply to other stages of pregnancy.

## METHODS

### Study approval and ethics statement

Samples obtained from non-pregnant (n=56) and pregnant women (n=58), with matched decidual single-cell suspensions (n=14), were used in this study (Supplementary Table 12). Samples from pregnant individuals used in LegendScreen and scRNAseq were at term gestation and >37 weeks. Samples from pregnant individuals used to measure concentrations of IL-6 and sIL-6R in blood plasma were all in the 3^rd^ trimester, ranging from 29-42 weeks (Supplementary Table 12). Experiments conformed to the Declaration of Helsinki Principles and the Australian National Health and Medical Research Council Code of Practice. Written informed consents were obtained from all participants prior to the study. The studies were approved by the Mercy Health (R14/25 and R04/29), Melbourne Health (HREC/66341/MH-2020), and University of Melbourne (#1443540, 2024-13344-58055-11, 2020-20782-12450-1) human research ethics committees (HRECS).

### Human blood processing

Whole blood was collected in heparinised blood tubes. Blood plasma was collected by centrifugation of whole blood at 300 *g* for 10 min. PBMCs were isolated through density-gradient centrifugation (Ficoll-Paque, GE Healthcare, Uppsala, Sweden). Isolated PBMCs were cryopreserved in fetal calf serum containing 10% DMSO.

### Human placenta tissue processing

Decidual tissue derived from placentae was processed as described previously^24^. Placental cotyledons were obtained from multiple locations on the maternal surface, before the basal plate and chorionic surface were removed and villous decidual tissue obtained from the middle cross-section. Tissue was washed in PBS and mechanically dissociated before enzymatic digestion in RPMI 1640 medium supplemented with 2 mg/mL Collagenase D (Roche, Basel, Switzerland), 0.2 mg/mL DNAase I (Roche), 100 U/mL penicillin, 100 µg/mL streptomycin and 10 mM HEPES. Digested tissue was put through a 70 μM filter before adding red blood cell lysis buffer (0.168 M NH_4_Cl, 0.01 mM EDTA, and 12 mM NaHCO_3_ in ddH_2_O). Single-cell suspensions of decidual tissue were cryopreserved in fetal calf serum with 10% DMSO.

### LegendSCREEN on CD4^+^ T cell subsets

Surface protein screening was performed using the Human LEGENDScreen PE kit (Biolegend, cat. no. 700011), as described previously^25^. In brief, PBMC or decidual single-cell suspensions were labelled with their respective CD45 and/or CTV fluorophore combinations to create sample barcodes. Sample were subsequently stained with the remaining backbone antibodies (Supplementary Table 13), before being multiplexed and distributed across the LEGENDScreen PE antibody wells. Samples were resuspended in MACS buffer and acquired on the Cytek Aurora 5-laser flow cytometer. Data were analysed in FlowJo v10 by first deconvoluting donor barcodes, followed by gating for CD4^+^ T cell subsets to determine frequencies of PE^+^ cells (Supplementary Figure 1A). Markers with >10% PE^+^ in at least one sample were included in analysis, and those with zero or near zero variance across all samples were removed.

### scRNAseq of PBMCs and decidual CD4^+^ T cells

scRNAseq was performed as described previously^25^. Briefly, PBMCs and decidua samples were stained with TruStain FcX (Biolegend, cat. no. 422301) for 10 min, followed by fluorescent antibody mix (Supplementary Table 14) for 30 min, and TotalSeq™-C Human Universal Cocktail (Biolegend, cat. no. 399905) and TotalSeq-C anti-human Hashtag antibodies 1-6 (cat. nos. 394661, 394663, 394665, 394667, 394669, 394671) for 30 min. Samples were washed and CD3^+^ T cells were sorted into RPMI 1640 with 10% fetal calf serum (Supplementary Figure 1B). Sorted T cells were multiplexed followed before performing scRNAseq according to 10x Genomics Next GEM Single Cell 5’ v2 manufacturer’s instructions using a Chromium iX.

Sequencing data were processed as described previously^25^. Briefly, raw data were processed with Cell Ranger v7.1.0 and samples demultiplexed based on TotalSeq-C Hashtag expression and differences in single nucleotide polymorphisms using cellsnp-lite v1.2.2 and vireo v0.2.3^26,27^. Downstream analyses was performed using Seurat v4.3.0.1^28^. The FindMarkers function from Seurat was used to identify differentially expressed genes and proteins. The gene *MTRNR2L8* had high expression in one non-pregnant individual and was not included in the interpretation of results. Gene set enrichment analysis was performed with clusterProfiler^29^, msigdbr, and enrichplot^30^ packages.

### Measurement of IL-6 in blood plasma

IL-6 cytokine levels were measured as part of the LEGENDplex™ Human Inflammation Panel 1 kit, per manufacturer’s instructions (BioLegend, San Diego, CA, USA). Samples were acquired on a BD CantoII, with BD FACS DIVA v8.0.1 software. Data were analyzed using LEGENDplex™ Data Analysis Software.

### Soluble IL-6 receptor ELISA

Soluble IL-6 receptor protein levels were measured with a DuoSet ELISA kit (R&D Systems, Minneapolis, MN, USA) according to the manufacturer’s instructions. In brief, 96-well Nunc Maxisorp ELISA plates (ThermoFisher, plasma) were coated overnight with the capture antibody overnight, before blocking with 1% w/v BSA for a minimum of 1□h. Blood plasma was diluted 1:300 and Standards were serial diluted before adding to the coated plate and incubating for 2□h at ambient temperature. Detection antibody was added for a further 2□h, and finally streptavidin-HRP, substrate solution and stop solution (2□N H_2_SO_4_) were added subsequently for 20□min each. A Multiskan plate reader (Labsystems) using the Thermo Ascent Software for Multiskan v2.4 was used to record optical densities at 450 and 540 nm. Data were interpolated to the standard curve using a 4-point logistic.

### Flow cytometry for validation of selected differentially expressed markers

Cryopreserved PBMCs or decidual single cell suspensions were thawed into pre-warmed RPMI containing 10% fetal calf serum, washed with MACS buffer (PBS containing 5% BSA) and subjected to surface antibodies (Supplementary Table 15) for 20 min at 37°C. Samples for intracellular staining were permeabilized in eBioscience Foxp3/Transcription Factor Staining Buffer kit (Thermo Fisher Scientific) before subjecting to intracellular antibodies (Supplementary Table 15). Samples were acquired on a BD LSRFortessa and analysed in FlowJo v10.

## Acknowledgements

We thank Brooke Henshall, Emily Allen, Kaitlin Constable, and Katelyn Dark for support with the pregnancy cohort, Jenny Anderson (Doherty Institute Computational Sciences Initiative), the Melbourne Cytometry Platform (University of Melbourne) and the Flow Cytometry and Cell Sorting Shared Resource (St Jude Children’s Research Hospital) for technical assistance. This research was funded in whole or part by the National Health and Medical Research Council (NHMRC) Investigator Grants: EL1 to LCR (#2026357) and THON (#1194036), EL2 to DAW (#1174555) and L2 to KK (#2033783). For the purposes of open access, the authors have applied a CC BY public copyright licence to any Author Accepted Manuscript version arising from this submission. JRH was supported by the Melbourne Research Scholarship from The University of Melbourne. This work was supported by MRFF Award (#2016062) to KK, PGT, THON and LCR, and NIAID UO1 grant 1U01AI144616-01 “Dissection of Influenza Vaccination and Infection for Childhood Immunity” (DIVINCI) to PGT and KK.

## Author Contributions

LCR and KK supervised the study. JRH, THON, LCR and KK designed the experiments. JRH, LCR, THON, EKA and MJS performed and analysed experiments. JRH, SL, JS and JCC analysed data. PGT provided crucial reagents. JAJ, SJK, KB, DAW, ML and SW recruited or provided patient cohorts. JRH, NdA, NJH, LCR and KK wrote the manuscript. All authors reviewed and approved the manuscript.

## FIGURE LEGENDS

**Supplementary Figure 1.**
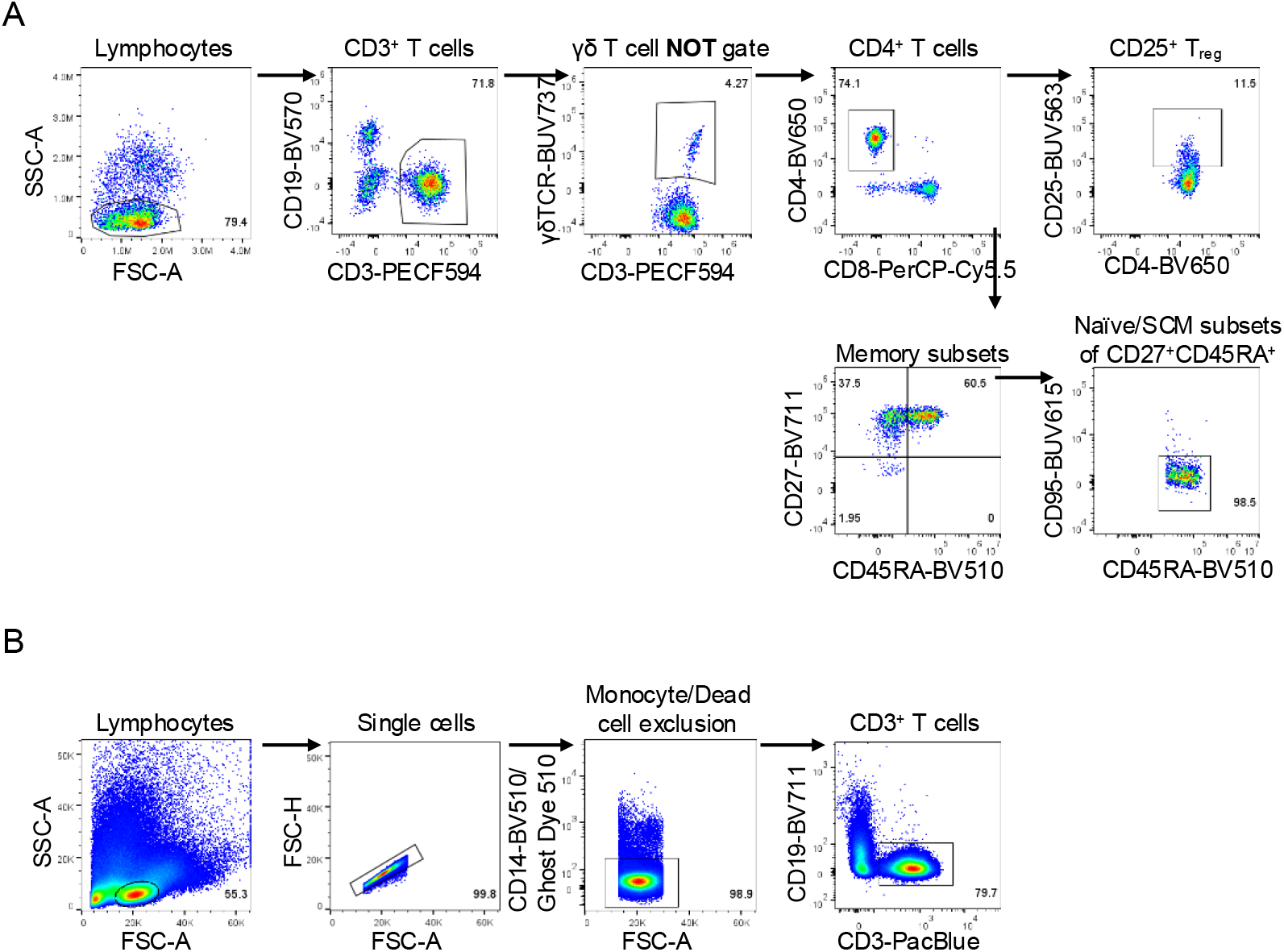
Gating strategies for LEGENDScreen and scRNAseq. (A) Gating strategy from pre-processed flow cytometry data as previously described^25^ to analyse T_Naive_, T_SCM,_ T_CM_, T_EM_, T_EMRA_, and T_reg_ CD4^+^ T cell subsets in LEGENDScreen. (B) Gating strategy to sort on T cells for single-cell RNA sequencing, of which CD4^+^ T cells were determined bioinformatically from RNA sequencing.

**Supplementary Figure 2.**
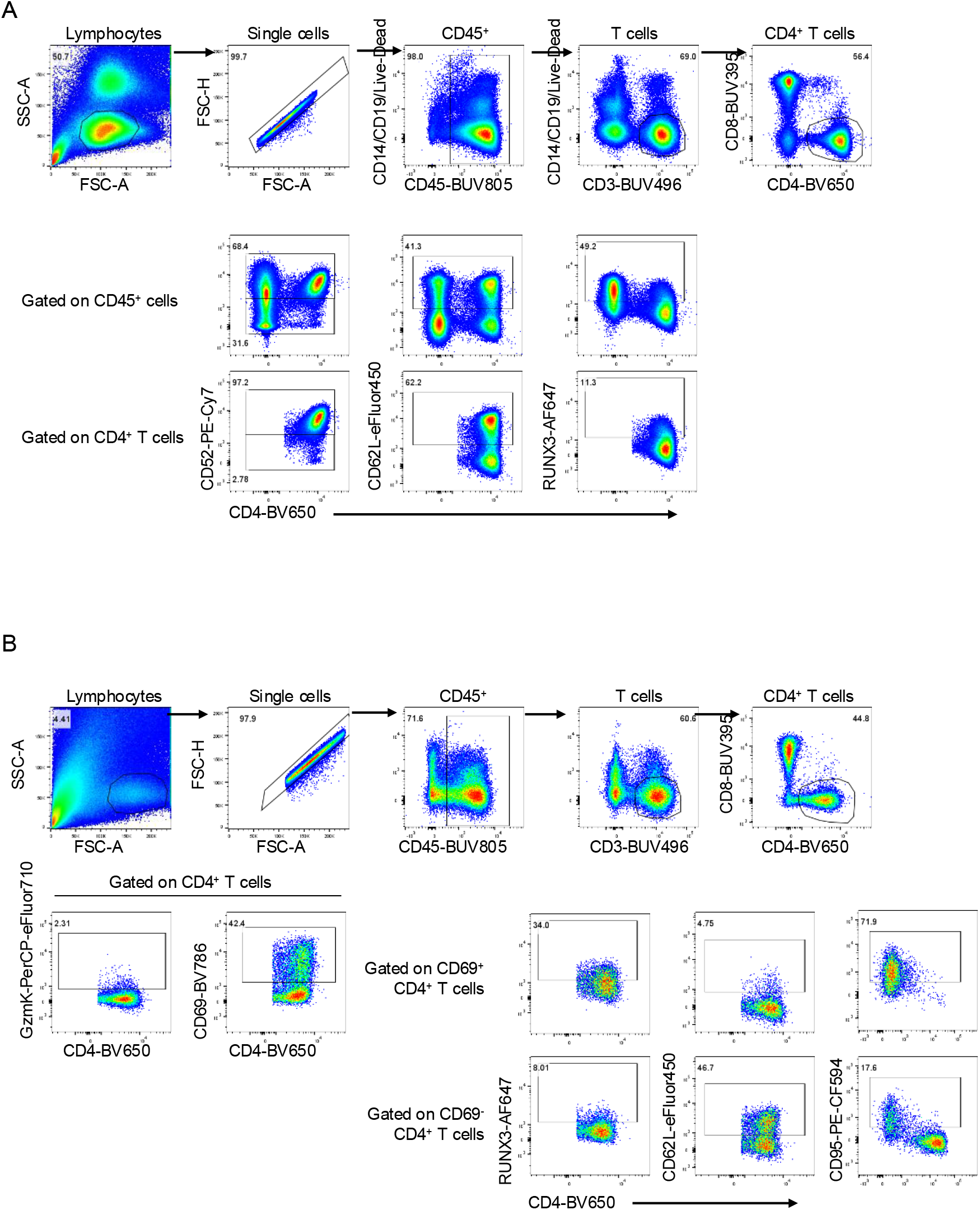
Gating strategy for validation flow cytometry. (A-B) Gating strategy to analyse selected differentially expressed markers on peripheral blood CD4^+^ T cells (A) and decidual CD4^+^ T cells (B). Gates were set on the CD45^+^ population to define thresholds before applying them to CD4^+^ T cells.

